# WRKY1 confers resistance to powdery mildew by accelerating SAR and preventing over-immunity in apple

**DOI:** 10.1101/2024.01.24.577112

**Authors:** Liming Lan, Lifang Cao, Lulu Zhang, Weihong Fu, Shenchun Qu, Sanhong Wang

## Abstract

Powdery mildew is one of the most serious diseases in apple production. SAR has a broad-spectrum immunity in plants against pathogen. Plants activate SAR against pathogen invasion and also prevent over-immunity. The relevant mechanism is still unknown in apple. In this study, we isolated and identified powdery mildew pathogen from the field and preserved them on the apple tissue culture seedlings. We performed DAP-seq of powdery mildew-inducible WRKY40. WRKY40 positively regulates NPR3like by directly binding to the W-box element of its promoter. NPR3like represses the expression of the PR1 gene in the presence of SA by competing with TGA2 for binding to NPR1. WRKY1 positively regulates WRKY40 by directly binding to the dual W-box element of its promoter, while WRKY1 positively regulates NPR3like by directly binding to the W-box element of its promoter. The expression trends of WRKY1, WRKY40, and NPR3like were basically the same as that of PR1 within 24 h after powdery mildew and SA treatments. Besides, WRKY1 increased SA content by positively regulating EPS1. After inoculation with powdery mildew, the up-regulation of PR1 in RNAi-silenced plants of WRKY1 was more slowly compared with the wild type, and the number of spores and mycelium increased significantly. In summary, we established a new model of NPR3like inhibition of NPR1 activity positively regulated by the WRKY1-WRKY40 module and found that the WRKY1-EPS1 module accelerated the up-regulation of PR1 by increasing the SA content. Finally, we elucidated WRKY1 confers resistance to powdery mildew by accelerating SAR and preventing over-immunity in apple.

## Introduction

Apple powdery mildew is one of the major diseases in apple production, a fungal disease caused by *Podospharea leucotricha* (Gañán et al. 2020), which mainly affects buds, leaves, new shoots, and young fruits of apples, and severely affects the tree’s vigor as well as the yield and quality of fruits (Zhang et al. 2021; Papp et al. 2016).

SA is an important plant immune signal for defense against pathogenic bacteria (An et al. 2011), and there are two different SA synthesis pathways in plants, namely the phenylalanine ammonia lyase (PAL) and the isobranchial acid synthase (ICS) pathways. About 10% of the pathogen-induced SA is from the PAL pathway, while about 90% is from the ICS pathway (Huang et al. 2010). In the ICS pathway, plastidic branch acids are converted to isobranchial acids by the isobranchial acid synthase ICS1/SID2 (Wildermuth et al. 2001), and transported to the cytoplasm by the MATE transporter protein EDS5 (Serrano et al. 2013), formation of glutamate-9-isobranchiate conjugate catalyzed by aminotransferase PBS3, and finally glutamic acid-9-isobranchial acid conjugate cleaves to form SA or in the presence of BAHD acyltransferase EPS1 (Rekhter et al. 2019; Torrens-Spence et al. 2019).

Systemic acquired resistance (SAR) is a defense mechanism induced in plants after infestation by pathogen and manifests itself as systemic broad-spectrum resistance (Muthamilarasan and Prasad 2013). SA is an important signaling substance to activate SAR (Horvath et al. 1996; Dong 2004; Uquillas 2004). Pathogen-induced SA breaks the disulfide bonds between NPR1 aggregates, and the SA-bound NPR1 monomer enters the nucleus and forms a protein complex with TGA2 to initiate the expression of PRs genes (Despres et al. 2000). NPR3 and NPR1 are genes belonging to the same class of NPRs and NPR3 plays a regulatory function by interacting with NPR1 protein (Wang et al. 2020). CUL3 (cullin 3) is an E3 ubiquitin ligase that ubiquitinates and degrades substrates (Li et al. 2021). When high levels of SA are present, Arabidopsis NPR3 acts as an adaptor between CUL3 and NPR1 to degrade NPR1 through ubiquitination and is a negative regulator of SAR (Backer et al. 2019; Fu et al. 2012).

WRKY transcription factors are involved in plant response to biotic and abiotic stresses (Pieterse et al. 2012). The plant WRKY transcription factor was discovered and named in 1994 (Ishiguro et al. 1994). The conserved WRKY structural domain consists of approximately 60 amino acid residues (Mahiwal et al. 2023). WRKY transcription factors activate or repress the transcription of multiple target genes by binding to specific DNA sequences (Yang et al. 2001). Many WRKY transcription factors in Arabidopsis, including: WRKY18, WRKY40, WRKY60, and WRKY70, increase defense against pathogen by participating in SA-mediated activation of SAR (Xu et al. 2006; Li et al. 2004). Arabidopsis WRKY28 acts as a positive regulator of ICS1 to regulate SA biosynthesis (Wang et al. 2015). Li et al. (2013) found that Arabidopsis WRKY70 and WRKY54 co-negatively regulate SA biosynthesis to participate in stress response. Knockdown of Arabidopsis WRKY33 resulted in pathogen-induced elevated SA levels (Rainer et al. 2012). Yu et al. (2001) found that SA-induced WRKY-binding proteins specifically recognize the W-box element of NPR1 and activate the expression of downstream PRs genes. Arabidopsis WRKY38 and WRKY62 act as negative regulators of plant basal defenses by repressing the expression of PR1 genes (Kim et al. 2008).

However, prolonged immunization activated by SA has an inhibitory effect on plant growth, and many Arabidopsis mutants exhibit high levels of SA accumulation accompanied by growth inhibition (Li et al. 2001; Kachroo et al. 2005). How to accelerate immunization while preventing the effects of over-immunity on one’s own growth and development is an important entry point for studying the balance between immunity and growth in plants. In this study, we isolated and characterized powdery mildew bacteria from the field and preserved them on the apple tissue culture seedlings, laying the foundation for research on the mechanism of resistance to powdery mildew. We identified NPR3like, a downstream gene of powdery mildew-induced WRKY40. NPR3like represses PR1 gene expression by competing with TGA2 for binding to NPR1 in the presence of SA, and there is no protein interaction between NPR3like and CUL3, suggesting that this is a novel mechanism for regulating the activity of NPR1 that is different from that of Arabidopsis NPR3. We identified WRKY1, an upstream regulatory gene of WRKY40, and at the same time, WRKY1 directly regulates the expression of NPR3like. The trends of WRKY1, WRKY40, and NPR3like expression after powdery mildew and SA treatments were essentially the same as those of PR1. In addition, WRKY1 directly positively regulated the SA synthesis-related gene EPS1. After inoculation with powdery mildew, the up-regulation of PR1 in RNAi-silenced plants of WRKY1 was more slowly compared with the wild type, and the number of spores and mycelium increased significantly. In summary, this study established a new model of NPR3like inhibition of NPR1 activity positively regulated by the WRKY1-WRKY40 module, found that the WRKY1-EPS1 module accelerated the up-regulation of PR1 by increasing the synthesis of SA, and elucidated WRKY1 confers resistance to powdery mildew by accelerating SAR and preventing over-immunity in apple, which lays a theoretical foundation for resolving the balance between plant immunity and growth.

## Results

### Isolation and characterization of powdery mildew pathogen

Powdery mildew is a biotrophic fungal pathogen, and isolating and purifying the pathogen is a challenging task that has hindered disease resistance research. To overcome this challenge, young leaves with powdery mildew were carefully selected from the field in spring when apples were just budding. Surface pathogen were removed from the spore-bearing leaves, and the leaf stems were placed in culture medium and incubated. After 7 days, re-growth of powdery mildew mycelia and spores on the leaf surface was observed (Fig. 1A). To ensure preservation without contamination by stray bacteria, the isolated pathogen was inoculated on rooted tissue culture seedlings (Fig. 1F). Visible spots on the leaves were observed after 8 days (Fig. 1B) and the pathogen proliferated after 40 days (Fig. 1C). The large number of ellipsoidal conidia observed during microscopic observation of the powdery mildew spores during reproduction (see Fig. 1, D and E) strongly suggests the successful isolation of promiscuous powdery mildew pathogen from the field. It also demonstrates the feasibility of preserving, propagating and inoculating powdery mildew pathogen on rooted tissue culture seedlings.

**Fig. 1.**
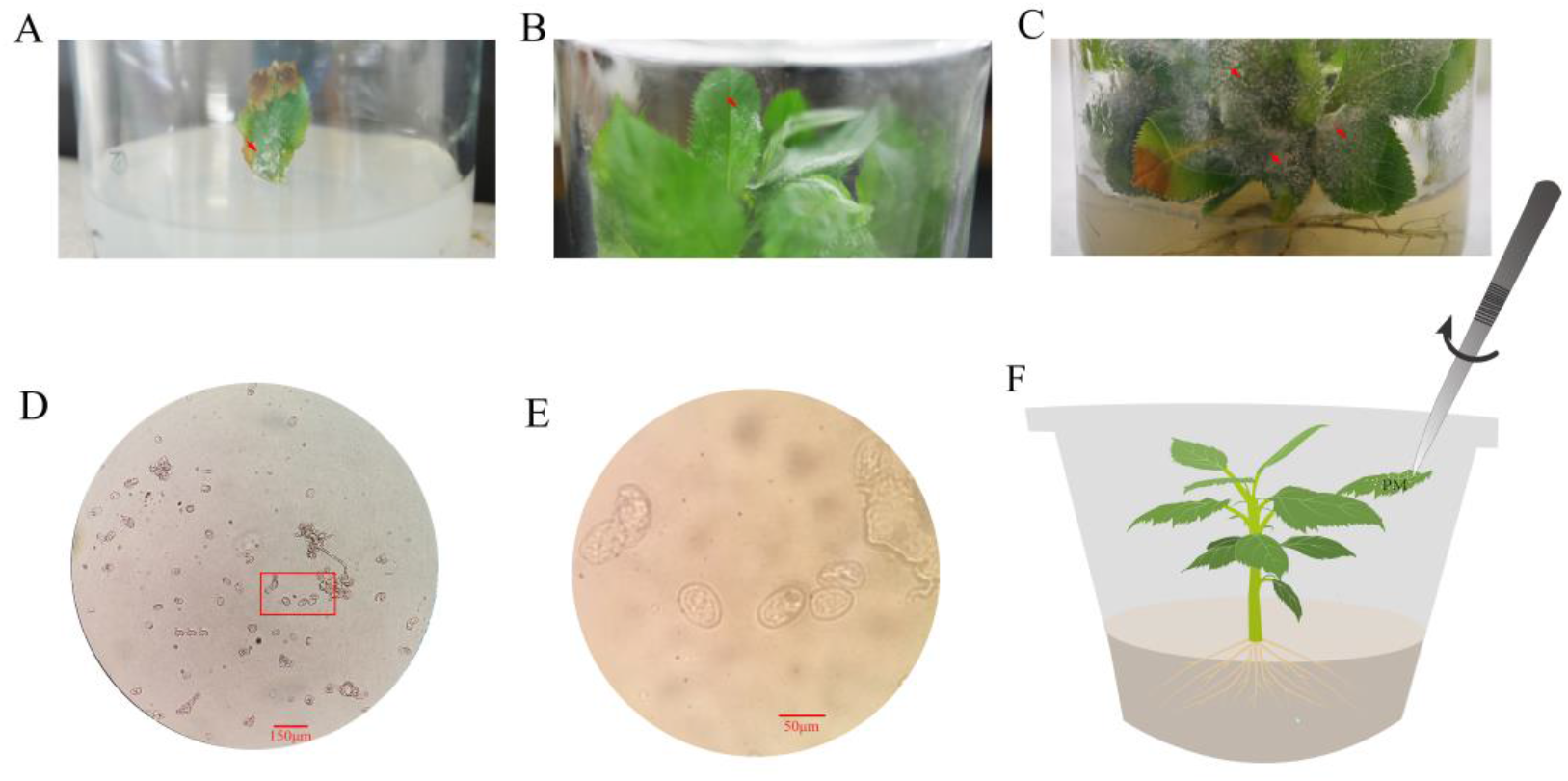
Isolation and identification of apple powdery mildew pathogen. A) Young field leaves carrying powdery mildew pathogen were removed from the surface of the disease, and the leaf stalks were inserted into the medium for 7 days. The leaves regrew mycelium and spores of powdery mildew pathogen. B) Isolated powdery mildew pathogen was inoculated on rooted tissue culture seedlings for 8 days before the appearance of visible spots on the leaf blades. C) The disease was proliferated after inoculation with powdery mildew pathogen for 40 days. D and E) Microscopic observation of powdery mildew spores using a light microscope. 20x microscopic observation (D), 100x microscopic observation (E) of the red boxed part. F) Schematic illustration of how powdery mildew pathogen are inoculated and preserved. Tweezers were used to clip the diseased leaves and rub them on the rooted tissue culture seedlings.

### Powdery mildew invasion promotes WRKY40 accumulation and SA-mediated SAR activation

We started from the six candidate WRKY genes with powdery mildew resistance obtained from a previous study (Lan et al. 2021), and the RT-qPCR results showed that all six WRKY genes were significantly up-regulated after inoculation with powdery mildew, among which WRKY40 was more significantly up-regulated compared with the other genes (Fig. 2A), and the protein accumulation and gene expression of WRKY40 after powdery mildew bacteria treatment showed a consistent trend (Fig. 2B). WRKY40 was localized in the nucleus (Supplemental Fig. S1). Therefore, the present study prioritized the analysis of the mechanism by which WRKY40 regulates powdery mildew resistance.

**Fig. 2.**
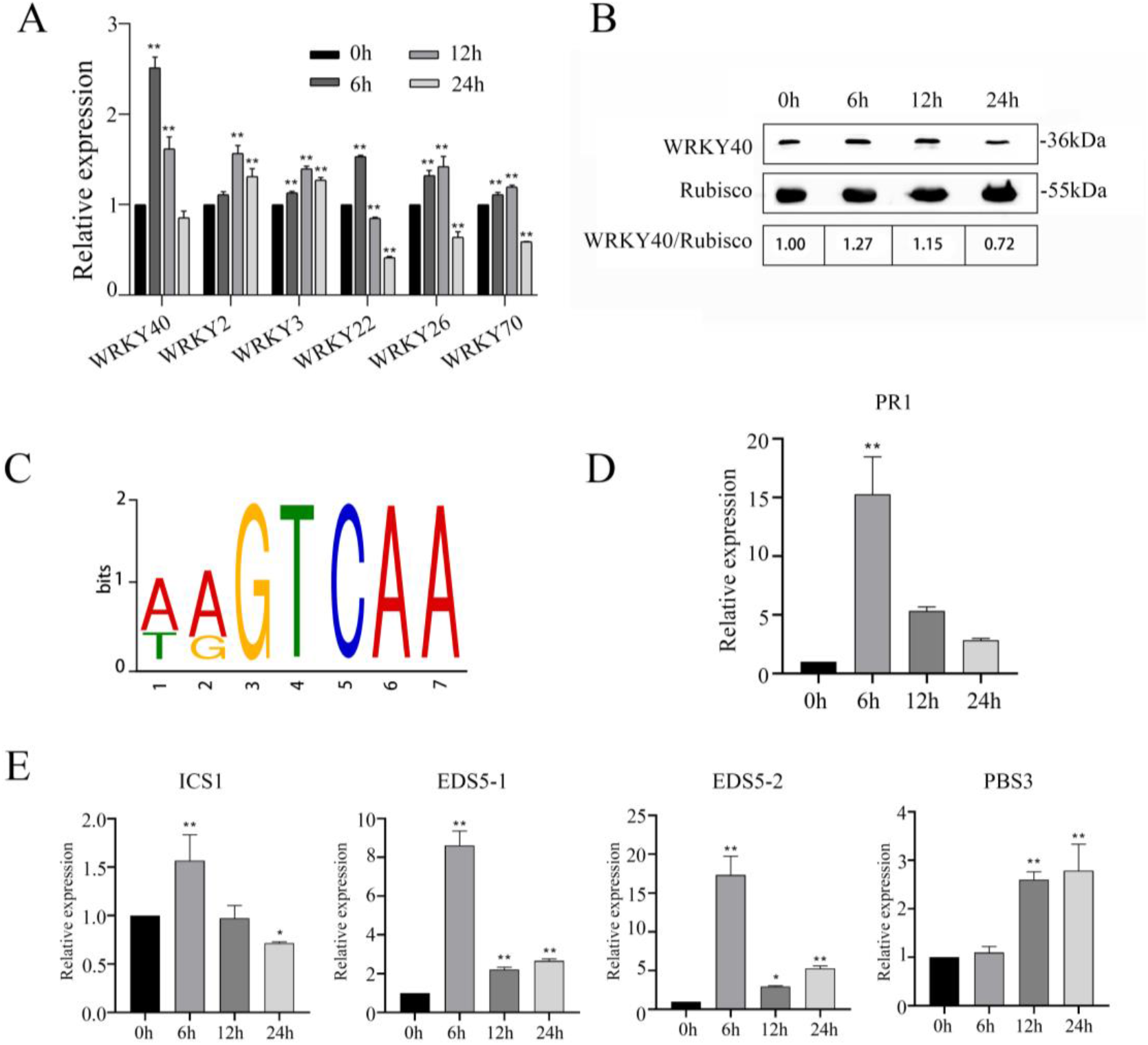
Powdery mildew invasion promotes WRKY40 accumulation and SA-mediated activation of SAR. A) Using wild-type plants that had been rooted for 40 days, RT-qPCR showed that expression of six WRKY genes candidate for powdery mildew resistance after inoculation with powdery mildew. B) Using wild-type plants that had been rooted for 40 days, after inoculation with powdery mildew, Western Blot was performed to detect WRKY40 protein accumulation. Rubisco was used as Loading control. WRKY40/Rubisco is the ratio of gray scale values of protein bands. C) DAP-seq results showed that WRKY40 binding sites were mainly enriched in the W-box (GTCAA). D) Using wild-type plants that had been rooted for 40 days, RT-qPCR showed that expression of the PR1 gene after inoculation with powdery mildew. E) Using wild-type plants that had been rooted for 40 days, RT-qPCR showed expression of four SA synthesis-related genes after inoculation with powdery mildew. The data represent the means and standard deviations of three independent replicate experiments. Asterisks (*) indicate significant differences from the control (Student’s t test, **P < 0.01).

In order to screen the candidate downstream genes of WRKY40, we performed DAP-seq on WRKY40, and the results showed that the major binding element of WRKY40 was the W-box (Fig. 2C). We performed KEGG on 380 candidate downstream genes of WRKY40, and the results showed that 185 genes were significantly enriched in “Metabolic pathways”, 28 genes were enriched in “Plant hormone signal transduction”, and 26 genes were enriched in “Plant-pathogen interaction” (Supplemental Fig. S2). This suggests that WRKY40 may be involved in the regulation of hormone signaling pathways.

SA is an important hormone regulating plant resistance. RT-qPCR results showed that the expression of PR1 gene was significantly up-regulated after inoculation with powdery mildew, which peaked at 6 h and then gradually decreased (Fig. 2D). The expression of several SA synthesis-related genes, ICS1, EDS5, and PBS3, was significantly up-regulated (Fig. 2E). It suggests that the invasion of powdery mildew pathogen promotes SA synthesis to activate SAR.

### WRKY40 specifically binds to W-box elements of NPR3like and positively regulates its expression

To analyze whether WRKY40 is involved in regulating SAR, we screened three NPRs genes from the DAP-seq results. Yeast one-hybrid assay showed that WRKY40 binds to the W-box element of the NPR3like gene, and mutation of the W-box element inhibited the binding of WRKY40, which is located at the upstream of the ATG at the position of -800 (Fig. 3, A and B). EMSA assay showed the same results as the yeast one-hybrid assay (Fig. 3C). Dual-luciferase reporter assays showed that WRKY40 positively regulated NPR3like expression, and mutations in the W-box element suppressed WRKY40 regulation (Fig. 3D). RNAi-silenced plants of WRKY40 were obtained, and RT-qPCR revealed that all three strains showed silencing effects (Fig. 3E). Ri-WRKY40#1 was selected for subsequent experiments, and RT-qPCR results showed that NPR3like expression was significantly suppressed in the silenced plants compared to the wild-type plants (Fig. 3F). It indicates that WRKY40 specifically binds to the W-box element of NPR3like to positively regulate its expression.

**Fig. 3.**
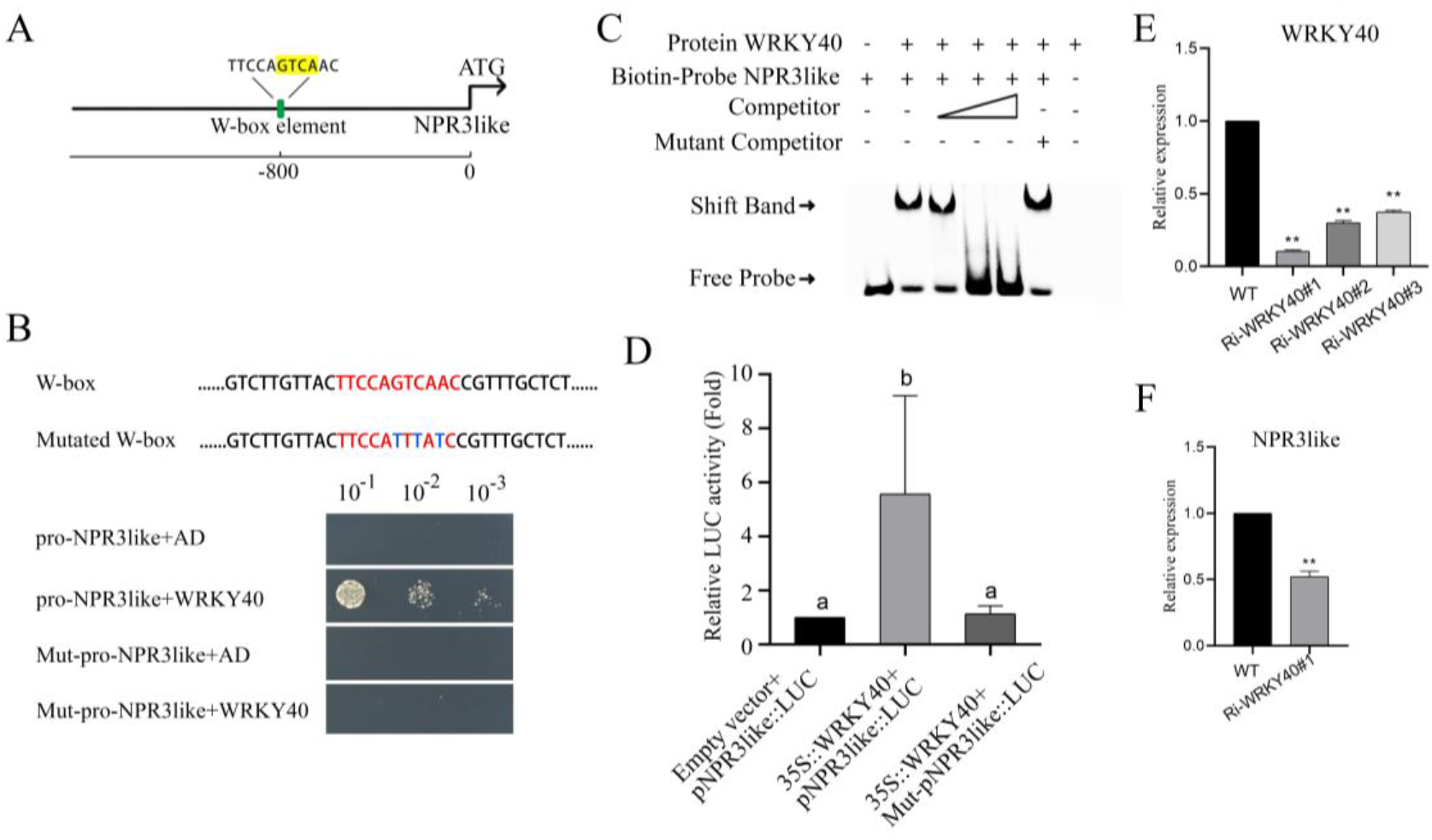
WRKY40 specifically binds to the W-box element of NPR3like and positively regulates its expression. A) Schematic representation of the position of the W-box element of the NPR3like promoter. B) Results of yeast one-hybrid assay for WRKY40 specifically binding to the W-box element of NPR3like. Yeast solution with OD of 0.45 was diluted to 10-1, 10-2, and 10-3 concentration gradients for validation. Sequences of binding elements are in red font and sequences after element point mutation are in blue font. C) Results of EMSA for WRKY40 specific binding to the W-box element of NPR3like. Biotin-labeled probes were selected to sequences of length 30 bp contain the W-box element, and the mutated competitor sequences are referenced in Figure B. D) Results of the dual-luciferase reporter assay for WRKY40 positively regulating NPR3like. The NPR3like promoter is sequence of length 2k upstream of the ATG. The mutated NPR3like promoter sequence is referred to Figure B. E, F) RT-qPCR analysis of WRKY40 (E) and NPR3like (F) expression in RNAi-silenced lines of WRKY40 using plants rooted for 20 days. The data represent the means and standard deviations of three independent replicate experiments. Asterisks (*) indicate significant differences from the control (Student’s t test, **P < 0.01).

### NPR3like represses PR1 expression by competing with TGA2 to bind NPR1 in the presence of SA

To explore whether NPR3like is involved in regulating NPR1 activity. The Arabidopsis NPR1 protein sequence was used as a template to compare to the apple genome to obtain all NPRs genes in apple (Supplemental Fig. S3, A and B), and the gene ID of apple NPR1 was determined by evolutionary tree analysis (Supplemental Fig. S3C). Yeast two-hybrid assay showed that NPR3like and NPR1 exhibited protein interactions only in the presence of SA (Fig. 4A).

**Fig. 4.**
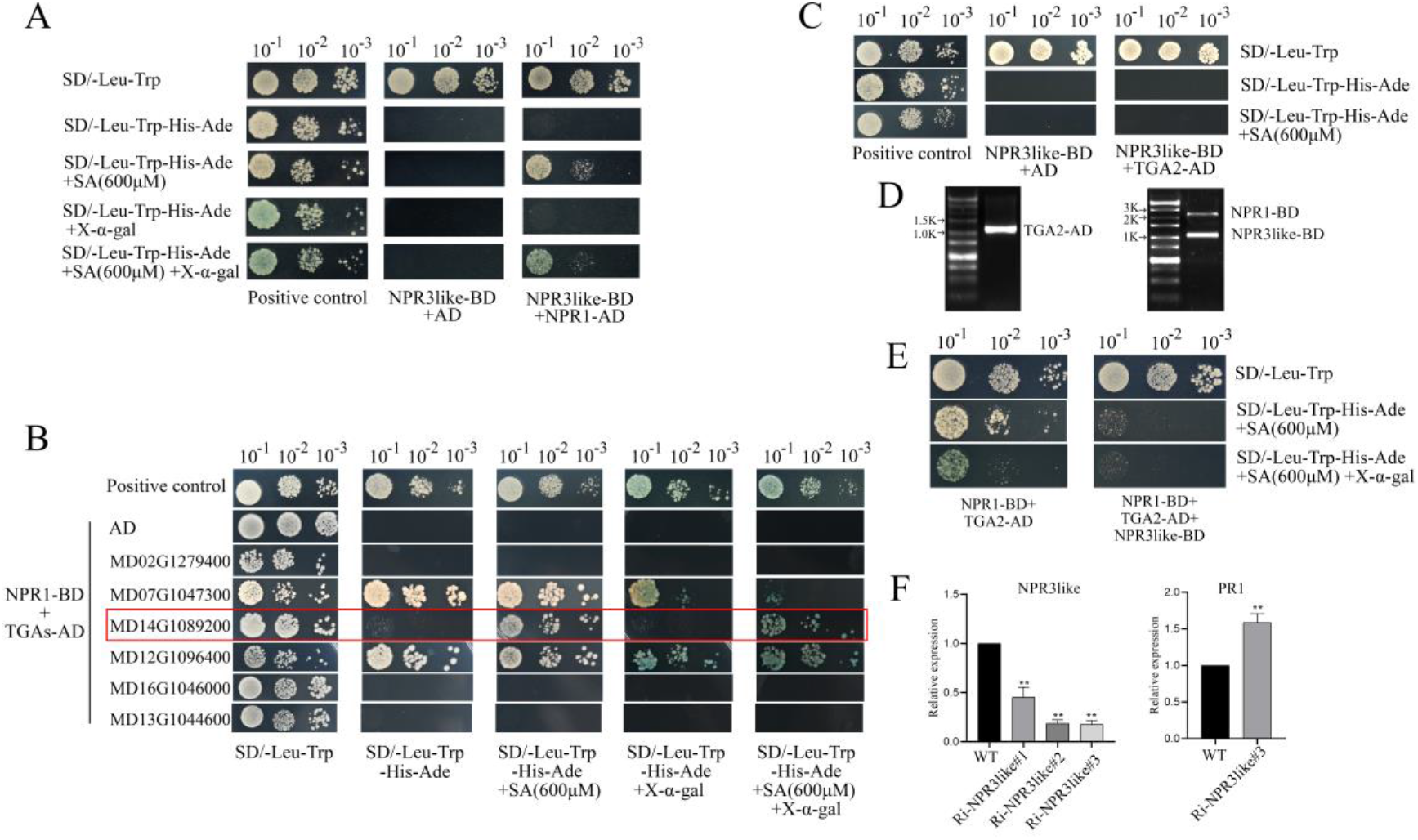
NPR3like represses PR1 expression by competing with TGA2 to bind NPR1 in the presence of SA. A) Yeast two-hybridization confirms protein interactions between NPR3like and NPR1 in the presence of SA. The concentration of SA in the medium was 600 μM. pGBKT7-53 and pGADT7-T were used as positive controls. Yeast solutions with an OD of 0.45 were diluted to 10-1, 10-2, and 10-3 concentration gradients for validation. B) Yeast two-hybrid results of multiple TGA proteins and NPR1 interactions validated. The red box shows the TGA protein that interacts with NPR1 in the presence of SA, which will be named TGA2. C) Yeast two-hybrid results of TGA2 and NPR3like interactions validated. D) Bacteriophage PCR validation of monoclonal bacteriophage after co-transfection of NPR1-BD, NPR3like-BD, and TGA2-AD plasmids into AH109 yeast receptor cell. E) Yeast three-hybrid results of NPR1, NPR3like, TGA2 three proteins interactions verified. F) Plants rooted for 20 days were used as material, RT-qPCR analysis the expression level of NPR3like and PR1 in RNAi silencing strain of NPR3like. The data represent the means and standard deviations of three independent replicate experiments. Asterisks (*) indicate significant differences from the control (Student’s t test, **P < 0.01).

Analysis of the conserved structural domains showed that the apple NPR3like protein consists of a BTB/POZ structural domain, and the Arabidopsis NPR3 protein contains BTB/POZ, DUF3420, Ank2, and NPR1like structural domains (Supplemental Fig. S4). Therefore, we hypothesized that the two have different functions. To confirm this speculation, we performed apple NPR3like and CUL3 interactions validation experiments. The Arabidopsis CUL3 protein sequence was used as a template to compare to the apple genome to obtain 9 CUL3 genes (Supplemental Fig. S5, A and B), and yeast two-hybrid assay showed that no protein interactions between apple NPR3like and any of the 9 CUL3s (Supplemental Fig. S5C). It suggests that apple NPR3like does not function as an adaptor between CUL3 and NPR1 to regulate NPR1 activity.

We hypothesized that NPR3like may affect the interaction of NPR1 with other proteins under SA. Arabidopsis TGA2, 5, and 6 participate in SAR by interacting with NPR1. The Arabidopsis TGA2 protein sequence was used as a template to compare to the apple genome to obtain all TGA genes in apple (Supplemental Fig. S6, A and B), and the gene IDs of apple TGA2, 5, and 6 were determined by evolutionary tree analysis (Supplemental Fig. S6C). Yeast two-hybrid assay revealed that there was one TGA protein that interacted with NPR1 only in the presence of SA (Fig. 4B), and this TGA protein had no interaction with NPR3like (Fig. 4C), which was named TGA2. Yeast three-hybrid assay showed that NPR3like inhibited the interactions between NPR1 and TGA2 in the presence of SA (Fig. 4, D and E). The silencing of the NPR3like gene was performed using RNAi technology, and positive plants were obtained as determined by the bluing of leaves after GUS staining (Supplemental Fig. S7). RT-qPCR revealed that all three strains showed silencing effects (Fig. 4F). Ri-NPR3like#3 was selected for the subsequent experiments. The RT-qPCR results showed up-regulation of the expression of PR1 in the silenced plants compared to the wild-type plants (Fig. 4F). It suggests that NPR3like represses PR1 expression by competing with TGA2 to bind NPR1 in the presence of SA.

### WRKY1 specifically binds WRKY40 and NPR3like W-box elements and positively regulates their expression

To further understand the WRKY40 regulatory mechanism, we analyzed the promoter of WRKY40 and found that there is a double W-box element (DWE element) at -350 upstream of the ATG (Fig. 5A). Studies have shown the existence of WRKY subfamilyⅠgene binding dual W-box elements (Liu et al. 2021). Based on this, we performed a screening assay for WRKY subfamilyⅠ gene-binding WRKY40 DWE elements using yeast one-hybrid (Qin et al. 2022). The results showed that WRKY1 binds to the DEW element of WRKY40, and that mutation of any of the W-boxes of the DWE element inhibits the binding of WRKY1 (Fig. 5B). The EMSA assay showed results consistent with the yeast one-hybrid (Fig. 5C). Dual-luciferase reporter assays showed that WRKY1 positively regulates WRKY40 expression and mutations in the DEW element inhibited WRKY1 regulation (Fig. 5D). Silencing of the WRKY1 gene was performed using RNAi technology, and positive plants were obtained as determined by the bluing of leaves after GUS staining (Supplemental Fig. S8). RT-qPCR revealed that all three lines showed silencing effects (Fig. 5E). Ri-WRKY1#3 was selected for subsequent experiments. RT-qPCR results showed that WRKY40 expression was significantly suppressed in the silenced plants compared to the wild-type plants (Fig. 5F). Subcellular localization showed that WRKY1 was localized in the nucleus (Supplemental Fig. S9). It suggests that WRKY1 binds the DWE element of WRKY40 and positively regulates its expression.

**Fig. 5.**
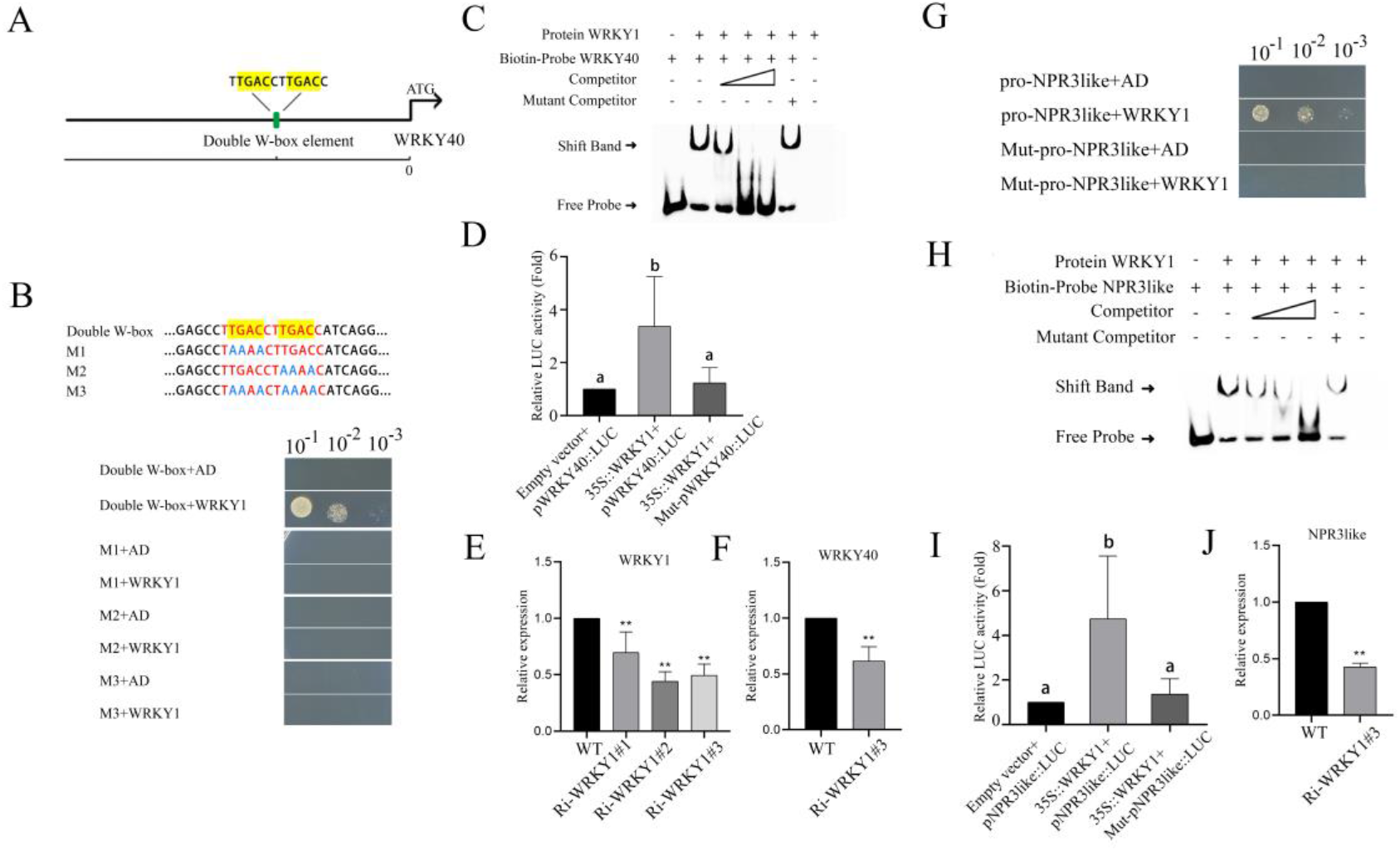
WRKY1 binds WRKY40 and NPR3like W-box elements and positively regulates their expression, respectively. A) Schematic representation of the DWE element position at the promoter of WRKY40. B) Results of yeast one-hybrid assay for WRKY1 binding to the DWE element of WRKY40. Yeast solutions with an OD of 0.45 were diluted to 10-1, 10-2, and 10-3 concentration gradients for validation. Sequences of the binding element are in red font and sequences after point mutation of the element are in blue font. C) EMSA results of WRKY1 binding to the DWE element of WRKY40. Biotin-labeled probes were selected to sequences of length 30 bp contain the DWE element, and the mutated competitor sequences are referenced to M3 in Figure B. D) Results of the dual-luciferase reporter assay for WRKY1-regulated WRKY40. The WRKY40 promoter is sequence of length 2k upstream of the ATG. The mutated WRKY40 promoter sequence is referenced to M3 in Figure B. E, F) Plants rooted for 20 days were used as material, RT-qPCR analysis of WRKY1 (E) and WRKY40 (F) expression in RNAi-silenced lines of WRKY1. G) Results of yeast one-hybrid assay for WRKY1 binding to the W-box element of NPR3like. Yeast solutions with an OD of 0.45 were diluted to 10-1, 10-2, and 10-3 concentration gradients for validation. Binding element sequences and mutated sequences are referenced in Figure 3, A and B. H) EMSA results of WRKY1-binding NPR3like W-box elements. Biotin-labeled probes were selected to sequences of length 30 bp contain the W-box element, and the mutated competitor sequence is referred to Figure 3B. J) Plants rooted for 20 days were used as material, RT-qPCR analysis of NPR3like expression in RNAi-silenced strains of WRKY1. The data represent the means and standard deviations of three independent replicate experiments. Asterisks (*) indicate significant differences from the control (Student’s t test, **P < 0.01).

Whether WRKY1 directly regulates the NPR3like gene. Yeast one-hybrid results showed that WRKY1 binds the W-box element of NPR3like, and mutations in the W-box inhibited WRKY1 binding (Fig. 5G). The EMSA assay showed results consistent with yeast one-hybrids (Fig. 5H). Dual-luciferase reporter assays showed that WRKY1 positively regulates NPR3like expression, and mutations in the W-box inhibited WRKY1 regulation (Fig. 5I). Ri-WRKY1#3 was selected for RT-qPCR assay, which showed that NPR3like expression was significantly suppressed in silenced plants compared with wild-type plants (Fig. 5J). It suggests that WRKY1 directly binds to the W-box element of NPR3like and positively regulates its expression.

### Powdery mildew- and SA-induced expression trends of WRKY1-WRKY40-NPR3like modules and PR1 were essentially identical

To analyze the expression trends of WRKY1-WRKY40-NPR3like module and PR1 after powdery mildew and SA induction. Wild-type apples were used as materials for inoculation with powdery mildew (Fig. 6A), the results showed that PR1 all showed significant up-regulation at 24 h after inoculation with powdery mildew, and then decreased after reaching a peak at 6 h. The expression trends of WRKY1 and WRKY40 were consistent with that of PR1. NPR3like all showed significant down-regulation, but the expression of NPR3like was significantly lower at 12 h and 24 h compared to 6 h (Fig. 6B).

**Fig. 6.**
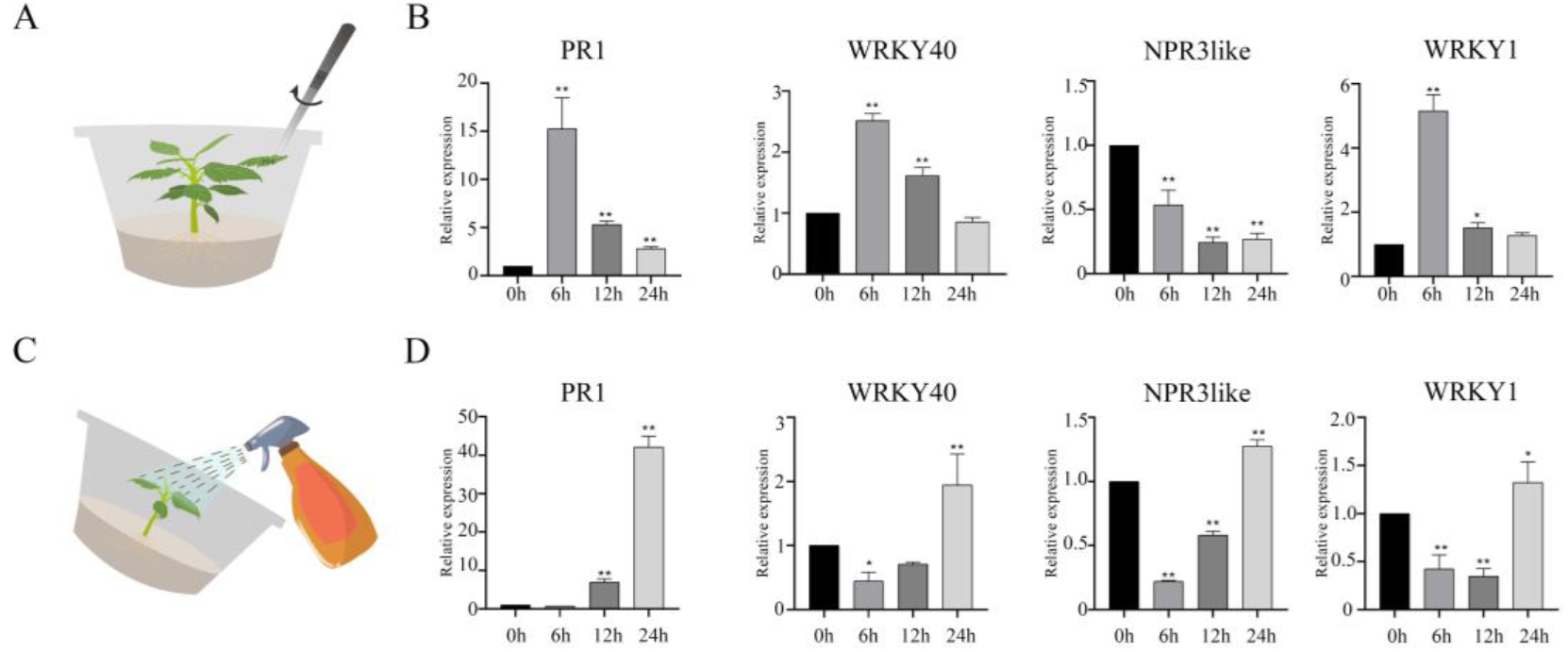
Expression of WRKY40, NPR3like, WRKY1, and PR1 after inoculation with powdery mildew and SA treatments, respectively. A) Schematic diagram of apple rooted tissue culture seedlings inoculated with powdery mildew. A diseased leaf cultured with powdery mildew for 30 d was selected and repeatedly rotated and rubbed between the leaves to be inoculated. B) Wild-type plants rooted for 40 days were used as material, RT-qPCR analysis of the expression of individual genes after inoculation with powdery mildew at 0 h, 6 h, 12 h, and 24 h. C) Schematic diagram of SA spraying on apple rooted tissue culture seedlings. 700 μL of SA at a concentration of 0.4 mM was sprayed. D) Wild-type plants rooted for 30 days were used as material, RT-qPCR analysis of the expression of individual genes at 0h, 6h, 12h, and 24h after spraying with SA. The data represent the means and standard deviations of three independent replicate experiments. Asterisks (*) indicate significant differences from the control (Student’s t test, **P < 0.01).

Next, we used wild-type apples as materials for spraying SA treatment (Fig. 6C). The results showed that PR1 expression showed a slow increase in 24h after SA treatment, and was up-regulated at 12h, and reached a higher level at 24h. The expression trends of WRKY1, WRKY40, and NPR3like were basically the same as those of PR1, with a slow increase at 6h, and up-regulation at 24h to reach a higher level (Fig. 6D). It suggests that the WRKY1-WRKY40-NPR3like module may be involved in regulating the rise and decrease of PR1 during the establishment of SAR in apple. Among other things, the mechanism by which the WRKY1-WRKY40-NPR3like module regulates PR1 lowering was identified in conjunction with the previous results. However, whether WRKY1 and WRKY40 are involved in the rise of PR1 is not known.

### WRKY1 directly binds to the W-box element of EPS1 to positively regulate its expression

To investigate whether WRKY1 and WRKY40 regulate the SA synthesis genes ICS1, PBS3, and EPS1 to participate in pathogen-induced SA synthesis. The Arabidopsis ICS1, PBS3, and EPS1 protein sequences were used as templates to compare to the apple genome to obtain all ICS1, PBS3, and EPS1 genes (Supplemental Fig. S10, S11, and S12). Yeast one-hybrid assay results showed that WRKY1 binds the W-box element of EPS1, and mutations in the W-box element inhibited WRKY1 binding, with the W-box element located upstream of the ATG at the position of-500 (Fig. 7A, B). The EMSA assay showed results consistent with the yeast one-hybrid assay (Fig. 7C). The dual-luciferase reporter assay showed that WRKY1 positively regulated EPS1 expression, and mutation of the W-box element suppressed WRKY1 regulation (Fig. 7D). Ri-WRKY1#3 was selected for RT-qPCR assay, which showed that EPS1 expression was significantly suppressed in silenced plants compared with wild-type plants (Fig. 7E). It suggests that WRKY1 directly binds to the W-box element of EPS1 and positively regulates its expression to participate in the up-regulation of PR1.

**Fig. 7.**
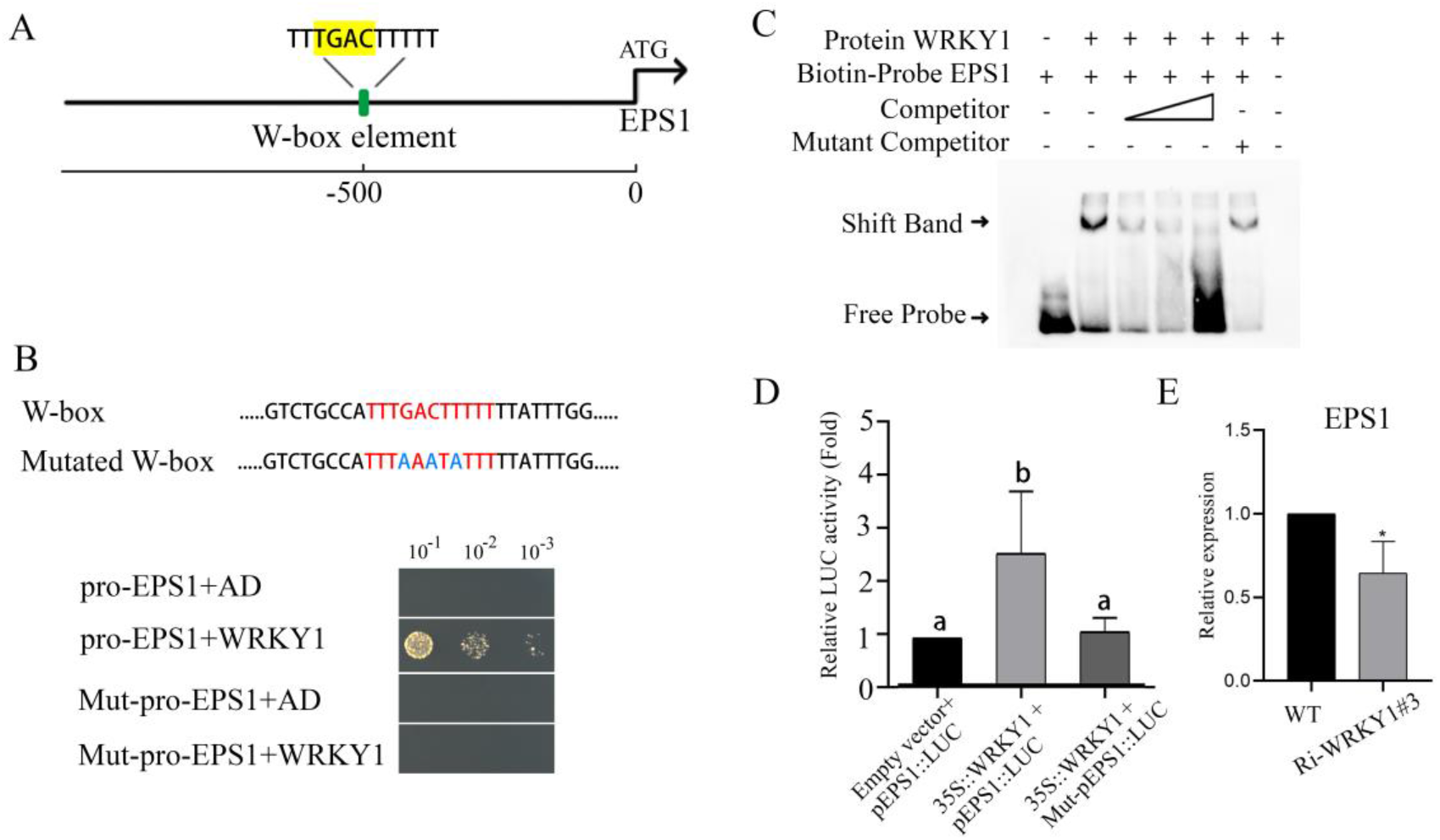
WRKY1 specifically binds to the W-box element of EPS1 and positively regulates its expression. A) Schematic representation of the position of the W-box element of the EPS1 promoter. B) Results of the yeast one-hybrid assay of the WRKY1-specific binding to the W-box element of EPS1. Yeast solutions with an OD of 0.45 were diluted to 10-1, 10-2, and 10-3 concentration gradients for validation. Sequences of binding elements are in red font and sequences after element point mutation are in blue font. C) EMSA results of WRKY1-specific binding to W-box elements of EPS1. The biotin-labeled probe was selected to sequence of length 30 bp contain the W-box element, and the mutated competitor sequences are referenced to Figure B. D) The dual-luciferase reporter assay for WRKY1-regulated EPS1. The EPS1 promoter is a sequence 2k upstream of ATG, and the mutated EPS1 promoter sequences are referenced to Figure B. E) Plants rooted for 20 days were used as material, RT-qPCR analysis the expression of EPS1 in RNAi silencing strains. The data represent the means and standard deviations of three independent replicate experiments. Asterisks (*) indicate significant differences from the control (Student’s t test, **P < 0.01).

**Fig. 8.**
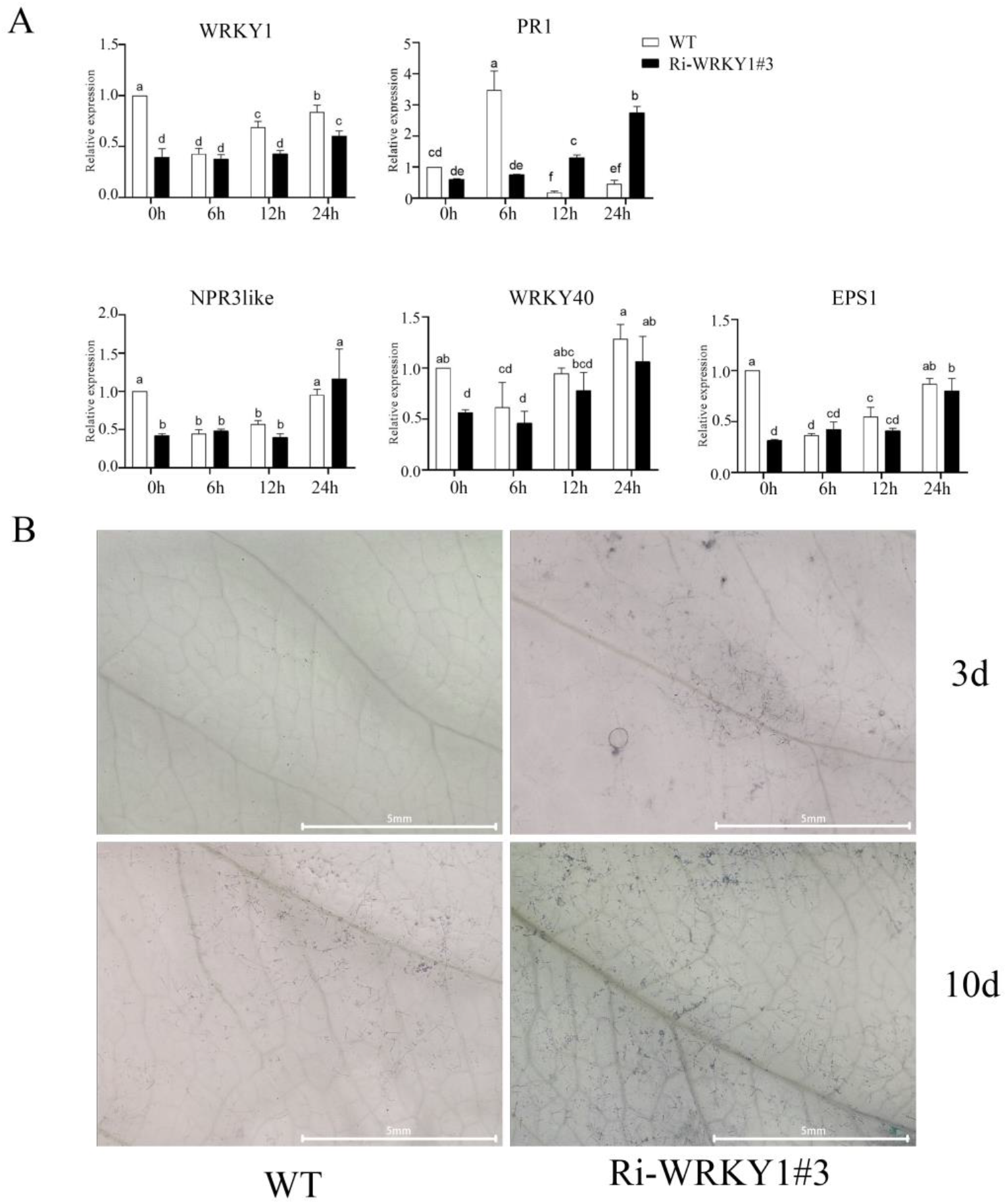
WRKY1-EPS1 module accelerates PR1 rise to increase powdery mildew resistance. A) Plants rooted for 20 days were used as material, the expression of WRKY1, PR1, WRKY40, NPR3like, and EPS1 gene after inoculation with powdery mildew in Ri-WRKY1-silenced plants. The data represent the means and standard deviations of three independent replicate experiments. Asterisks (*) indicate significant differences from the control (Student’s t test, **P < 0.01). B) Ri-WRKY1-silenced plants with 20 days of rooting were inoculated with powdery mildew at 3 and 10 days, and were stained using trypan blue staining solution for spores and hyphae. After decolorization, observations were recorded using a super field depth stereomicroscope with a magnification of 64×. The scale is 5 mm.

**Fig. 9.**
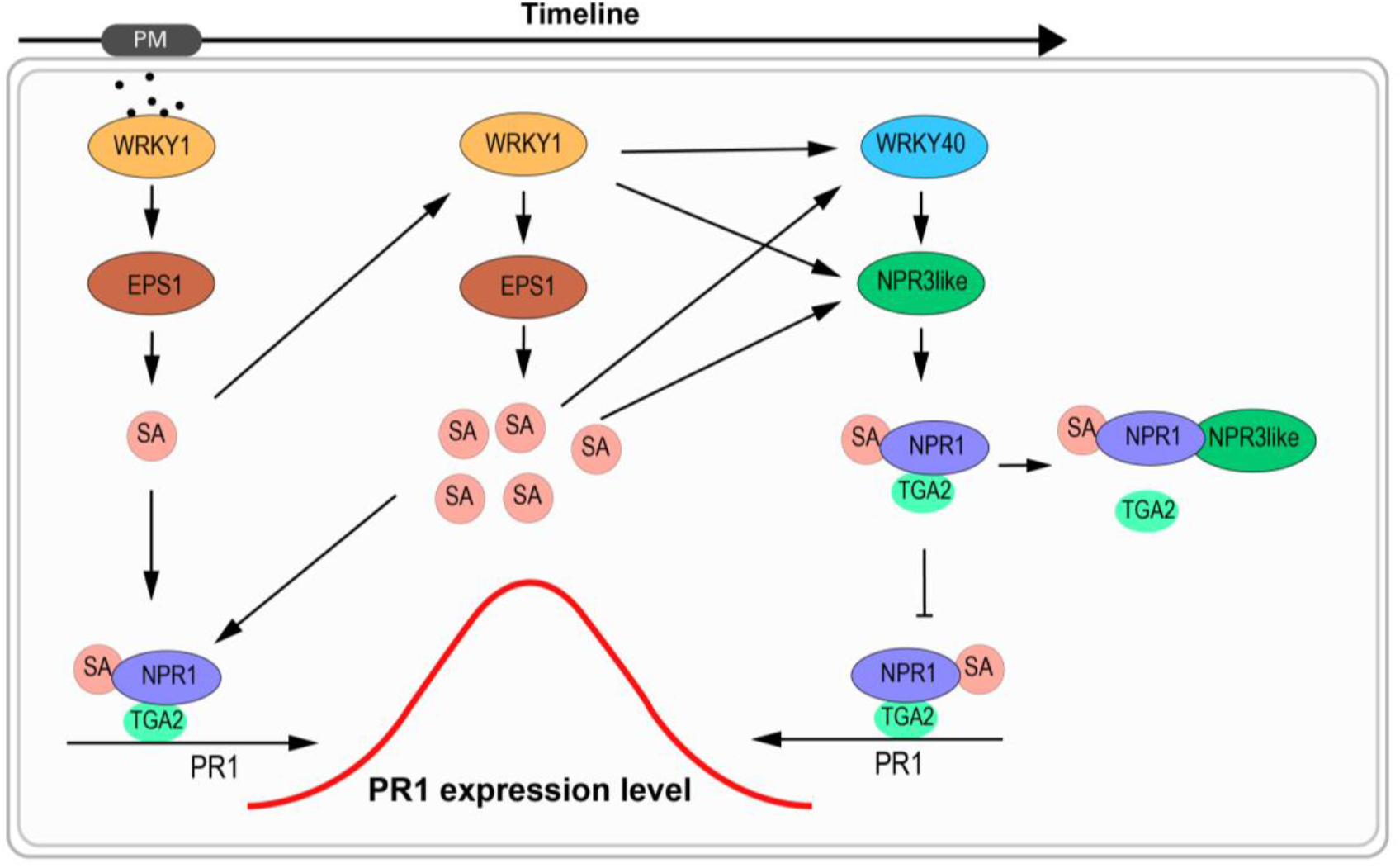
The model of WRKY1 confers resistance to powdery mildew by accelerating SAR and preventing over-immunity in apple. In response to powdery mildew invasion, plants produce signaling substances, and WRKY1 senses the signaling substances and synthesizes SA by activating the EPS1 gene. SA breaks the disulfide bond between the NPR1 polymers, and the NPR1 monomers that have bound SA enter the nucleus and form a protein complex with TGA2, which binds to the as-1 motifs of PR1 and initiates the expression of PR1 to defend against the invasion of powdery mildew. Meanwhile, SA would further stimulate the WRKY1-EPS1 model to produce more SA and accelerate the rise of PR1. With the accumulation of SA, the expression of the WRKY1-WRKY40-NPR3like model gradually increased, and NPR3like inhibited PR1 expression by competing with TGA2 to bind NPR1, preventing the effect of over-immunity on the growth and development of the plant itself.

### WRKY1-EPS1 module accelerates PR1 rise to increase powdery mildew resistance

To analyze the function of WRKY1-EPS1 module in regulating the rise of PR1. RNAi plants of WRKY1 were used as materials for powdery mildew inoculation, and the results showed that the expression of WRKY1 in RNAi silenced plants was significantly lower than that of wild-type plants within 24h of inoculation, and the rise of PR1 was more slow compared to wild-type. PR1 in wild-type plants reached the peak at 6h and then began to decline, while PR1 in silenced plants began to be up-regulated at 12h and reached a higher value at 24h. The expression of NPR3like, WRKY40 and EPS1 in WRKY1-silenced plants was significantly lower than that in wild-type plants within 24 h of inoculation, and the expression trend was basically consistent with that of WRKY1.

At 3 and 10 days after inoculation, leaves were stained with trypan blue staining solution to observe the number of spores and mycelium. The results showed that at 3 days, there was no obvious mycelium in the wild type, and a small number of spores and mycelium appeared in the silenced plants, and at 10 days, a large number of spores and mycelium appeared in both materials, but the number of spores and mycelium in the silenced plants was significantly higher than that in the wild type. Taken together, this suggests that the WRKY1-EPS1 module improves powdery mildew resistance by accelerating the rise of PR1.

## Discussion

Apple powdery mildew is a biotrophic fungal pathogen, and the acquisition and preservation of pathogen is one of the main problems that hinder research on apple powdery mildew resistance. Outbreaks of apple powdery mildew fungus occur mainly during the budding and fall growth periods, and we chose to isolate the powdery mildew fungus from the field during the spring budding period when the level of other pathogen in the field was low. After we obtained the isolated powdery mildew pathogen, we succeeded in preserving and propagating the pathogen on rooted tissue culture seedlings for the first time. This preservation guaranteed that the pathogen would not be contaminated by other stray pathogen again during propagation, which laid the foundation for subsequent resistance studies in this study, and also provided ideas for powdery mildew preservation studies in other species.

The WRKY family plays an important regulatory role in plant response to biotic and abiotic stresses (Pieterse et al. 2012). In this study, we identified WRKY40 genes in response to powdery mildew invasion and screened candidate downstream genes of WRKY40 using DAP-seq technology. These candidate genes were mainly enriched in hormone-related metabolic pathways, and we hypothesized that WRKY40 is involved in hormone signaling pathways to regulate powdery mildew resistance. SA-induced activation of SAR is an important preventive tool against disease invasion in plants (Muthamilarasan and Prasad 2013), and we found that apple powdery mildew invasion stimulates the synthesis of SA and PR1 significant up-regulation. Based on this, our screen of WRKY40 downstream genes focused on genes involved in the regulation of SAR. We found that WRKY40 directly regulates NPR3like expression.

In Arabidopsis, NPR1 acts as a receptor for SA to initiate PR1 expression, and NPR3 and NPR4 function by regulating NPR1 activity through interactions with NPR1, which acts as adaptors between CUL3 and NPR1 to ubiquitinate degradation of NPR1 (Fu et al. 2012). We found that apple NPR3like interacts with NPR1 in the presence of SA. The apple NPR3like protein consists of a BTB/POZ structural domain, and the Arabidopsis NPR3 protein contains BTB/POZ, DUF3420, Ank2, and NPR1like structural domains (Supplemental Fig. S4), and, second, the apple NPR3like has no protein interaction with CUL3 (Supplemental Fig. S5). It indicates that the mechanism by which apple NPR3like regulates NPR1 is different from that of Arabidopsis. In Arabidopsis, the NPR1 monomer that binds SA enters the nucleus and forms a protein complex with TGA2 to initiate PR1 expression (Despres et al. 2000). We found that there is a TGA2 protein in apple that interacts with NPR1 only in the presence of SA. Apple NPR3like competes with TGA2 protein for binding to NPR1 in the presence of SA to inhibit PR1 expression, and PR1 expression was significantly increased in NPR3like-silenced plants compared with the wild type. This suggests that we have discovered a novel mechanism of NPR3like regulation of NPR1 activity that is different from that of Arabidopsis.

We performed a preliminary analysis of the binding sites of apple NPR3like and NPR1 protein interactions. In Arabidopsis, the zinc finger structure of BTB/POZ of NPR1 binds Ank4 to stabilize NPR1, and SA unbinds the zinc finger structure from Ank4 to activate NPR1 (Wang et al. 2020). Apple NPR3like consists of a BTB/POZ structural domain, and apple NPR1 is consistent with the Arabidopsis NPR1 structural domain (Supplemental Fig. S3 and S4). Therefore, we hypothesized that the zinc finger structure of apple NPR3like binds Ank4 of NPR1.

Then, we performed a preliminary analysis of the binding sites of apple TGA2 and NPR1 protein interactions. We found that, in the absence of SA, apple NPR3like promotes TGA2 interactions with NPR1 (Supplemental Fig. S13), suggesting that, the zinc finger of the BTB/POZ structural domain is required for the binding of TGA2 to NPR1. Apple NPR3like does not interact with TGA2 (Fig. 4C), suggesting that only the zinc finger is insufficient for TGA2 binding to NPR1 and that other SA-unaffected structures of NPR1 need to be involved. In Arabidopsis, the binding site where TGA3 interacts with NPR1 is Ank1. therefore, we hypothesized that apple TGA2 binds both the zinc finger and Ank1 of NPR1.

The double W-box element is composed of two W-box elements of 12 bases that are specifically bound by WRKY family genes (Liu et al. 2021). We found a double W-box element in the WRKY40 promoter region, and the screening results showed that WRKY1 binds to the double W-box element of WRKY40 to positively regulate its expression, and at the same time, WRKY1 also directly positively regulates NPR3like. Taken together, we established a new model for the positively regulated NPR3like inhibition of NPR1 activity by WRKY1-WRKY40 module, and speculated that the purpose of this mechanism is to prevent the effect of over-immunity on self-growth and development. The results of powdery mildew and SA treatments showed that the expression trends of the WRKY1-WRKY40-NPR3like module and PR1 were basically the same, suggesting that the WRKY1-WRKY40-NPR3like module may be involved in the regulation of the rise and fall of PR1 during the establishment of SAR in apple. This result, on the one hand, provides support for the new model we established, and on the other hand, suggests a new idea: whether WRKY1 and WRKY40 are involved in the rise of PR1.

Pathogen-induced endogenous SA synthesis is an important factor affecting the expression trend of PR1 (An et al. 2011), and ICS1, PBS3, and EPS1 are the three major genes in the SA synthesis pathway (Huang et al. 2010). After screening, it was found that WRKY1 directly positively regulates EPS1 to promote SA synthesis. After inoculation with powdery mildew, the rate of PR1 rise in WRKY1-silenced plants was significantly reduced and the number of spores and hyphae was significantly increased compared with the wild type. It suggests that the WRKY1-EPS1 module improves powdery mildew resistance by accelerating the rise of PR1. In summary, this study established a new model of NPR3like inhibition of NPR1 activity positively regulated by the WRKY1-WRKY40 module, found that the WRKY1-EPS1 module accelerated the rise of PR1 by increasing SA synthesis, elucidated WRKY1 confers resistance to powdery mildew by accelerating SAR and preventing over-immunity in apple, provided a new idea for the study of the balance between immunity and growth and development.

## Materials and methods

### Isolation and characterization of powdery mildew pathogen

Young leaves carrying powdery mildew pathogen were taken from the field, leaving 0.5 cm long petioles, rinsed twice with tap water, soaked in 75% ethanol for 10 s, rinsed twice with sterile water, and soaked in 7% sodium hypochlorite for 2 min, rinsed twice with sterile water. The petioles were inserted into the medium to be cultured in the culture chamber. The medium formulation: 5 g/L agar + 50 mg/L benzimidazole (Deng et al. 2015). Leaves carrying powdery mildew fungus were picked up using tweezers and rubbed repeatedly on in apple rooted tissue culture seedlings for inoculation, preservation and propagation of the pathogen. Apple rooting medium formulation: 1/2 MS + 0.4 mg/L IBA + 1 mg/L IAA + 6 g/L agar + 30 g/L sucrose. Culture conditions in the incubation room: temperature of 25 degrees Celsius, photoperiod of 16h light and 8h dark.

### Powdery mildew infestation and SA spray treatments

To minimize other influencing factors, both powdery mildew and SA treatments were performed by selecting robust rooted tissue culture seedlings. Robustly grown rooted tissue culture seedlings were selected, and diseased leaves one month after powdery mildew infestation were clamped with forceps and rubbed repeatedly on apple leaves to complete the infestation. Robustly grown rooted tissue culture seedlings were selected and pruned to leave the top three fully expanded leaves that did not cover each other, so that the spray could fall on each part of the leaves, and 700 μL of SA solution at a concentration of 0.4 mM was sprayed. Samples were taken at 0, 6 h, 12 h, and 24 h after treatment and stored in a -80 refrigerator after liquid nitrogen flash freezing.

### Western Blot

In order to analyze the protein accumulation of WRKY40 after the invasion of powdery mildew, we used Western Blot to detect WRKY40. Antibody preparation for WRKY40 was done by Zoonbio Biotechnology (Song et al. 2021), and the Rubisco antibody was purchased from Sangon Biotech. The total protein was extracted from apple leaves at 0, 6, 12 and 24 h after inoculation with powdery mildew and subjected to Western Blot, using a concentration of 1:200 for WRKY40 and 1:2000 for Rubisco, and the grayscale values of the protein bands were calculated using Image J (Carvajal-Vergara et al. 2010).

### DAP-seq

To screen the downstream genes of WRKY40, DAP-seq analysis of WRKY40 was performed. We entrusted Bluescape Scientific to conduct this test. The CDS sequence encoding WRKY40 was constructed into a vector containing an affinity tag, a protein expression vector was constructed, and in vitro protein expression was carried out to form a fusion protein of WRKY40 and the affinity tag. Genomic DNA from apple leaves was extracted and a DNA library was constructed, then the in vitro expressed WRKY40 with affinity tag and the DNA library were combined, and subsequently the combined DNA was eluted and up-sequenced, and the obtained candidate downstream genes were subjected to KEGG clustering analysis (Bartlett et al. 2017).

### Yeast one-hybrid assays

The promoter fragment was constructed into pAbAi vector using the ClonExpress II One Step Cloning Kit (Item No. C112-01) from Vazyme, the recombinant pAbAi vector was linearized using BstB1 single enzyme digestion, and the digest product was transformed into Y1H yeast receptor cell, coated in SD/-ura medium, and positive monoclonal bacterial fluids were obtained for the self-activation test. The cds sequence of the transcription factor was constructed into the pGADT7 vector, and the recombinant pGADT7 plasmid transformed the yeast receptor cell containing the pAbAi vector, coated in SD/-Leu medium, and the pGADT7 empty load was used as a negative control, and positive monoclonal bacterial fluids were obtained for verification.

### EMSA

The CDS sequence encoding WRKY40 was constructed into the prokaryotic expression vector pCold-TF, and the recombinant pCold-TF plasmid was transformed into BL21 receptor cell for in vitro protein expression to obtain the purified WRKY40 protein. A 30 bp sequence containing the binding element was selected as a probe for biotin labeling. The probe without biotin labeling was used as a competitor. The probe with the binding element mutated was used as a competitor for the mutation.

### Dual-luciferase reporter

The CDS sequences of the transcription factors were constructed in pCAMBIA2300 overexpression vector (Hou et al. 2021), and the portion of 2k upstream of ATG was inserted into pLUC vector as the promoter sequence, and the point mutation of promoter sequences of the recombinant pLUC plasmid was accomplished by using the Mut Express II Fast Mutagenesis Kit V2 (Item No. C214-01) from Vazyme. The pCAMBIA2300 recombinant plasmid transformed the EHA105 receptor cell, and the pLUC recombinant plasmid transformed the EHA105 (pSoup) receptor cell. Summer is the rapid growth period of tobacco, at this time, 5 fully expanded leaves of robust tobacco were selected for the co-transformation test, the photoperiod was 16h light and 8h dark, 30 degrees incubated for 2 days, and then Dual Luciferase Reporter Assay Kit (Item No. DL101-01) from Vazyme was selected for the detection of LUC and Ren value. pCAMBIA2300 was used as a control in the empty load.

### Yeast two-hybrid assays

The CDS sequences of either of the two genes to be verified were constructed in the pGBKT7 vector, and the recombinant pGBKT7 plasmid and the pGADT7 empty plasmid were co-transformed into the AH109 receptor cell, coated in SD/-Leu-Trp medium, and after obtaining a positive clone, the autoreactivation was verified in the SD/-Leu-Trp-Ade-His medium. The CDS sequence of another gene was constructed in pGADT7 vector, pGBKT7 recombinant plasmid and pGADT7 recombinant plasmid were co-transformed into AH109 receptor cell, coated in SD/-Leu-Trp medium, and after obtaining a positive clone, the presence of protein interactions was verified in SD/-Leu-Trp-Ade-His medium, the concentration of SA in the medium was 600 μM and the concentration of X-α-gal was 20 mg/L. pGBKT7-53 and pGADT7-T were used as positive controls (Zhao et al. 2017).

### Yeast three-hybrid assays

To verify how NPR3like affects protein interactions of NPR1 and TGA2, we performed a yeast three-hybrid assay (Glass et al. 2015). The cds sequence of NPR1 was constructed on pGBKT7 vector, the cds sequence of NPR3like was also constructed on pGBKT7 vector, and the cds sequence of TGA2 was constructed on pGADT7 vector. The three recombinant plasmids were co-transformed into AH109 receptor cell, coated with SD/-Leu-Trp medium, and multiple single clones were picked up, and simultaneous presence of two pGBKT7 recombinant plasmids in a single clone was ensured by bacteriophage PCR. Positive clones were selected for validation in SD/-Leu-Trp-Ade-His medium. Single clones after co-transformation of NPR1 and TGA2 recombinant plasmids were used as controls. The concentration of SA in the medium was 600 μM and the concentration of X-α-gal was 20 mg/L.

### RNAi

After identifying the WRKY40-specific 500 bp cds fragment, this specific fragment and its complementary sequence were constructed on the plant expression vector pB7GWIWG2(II) using Gateway technology, and glufosinate ammonium was used as a screening marker. With the popularization and application of homologous recombination technology, we used a simpler method to construct RNAi vectors. The pRNAi-E vector was obtained by inserting the intron sequence derived from the pKANNIBAL vector at the polyclonal site of the pCAMBIA2300 vector. On this basis, the RNAi vector for the target gene can be constructed by ligating the forward and reverse fragments of the specific sequence of the target gene to both sides of the intron respectively (Song et al. 2017). According to this approach, we successfully constructed RNAi vectors for WRKY1 and NPR3like, and Kana was used as a screening marker. pCAMBIA2300 vector carries GUS tags, which can be used to verify whether the transgenic plants are positive or not by means of GUS stain. The leaves of the obtained transgenic plants were immersed in GUS staining solution, incubated at 37 overnight, and observed after destained by 95% alcohol.

### RT-qPCR

RT-qPCR experiments were done using QuantStudio 5 instrument. All gene IDs and primers for this study are shown in Supplemental Table S1.

### Trypan Blue Stain

The leaf samples were placed in trypan blue staining solution in a boiling water bath for 2 min and left at room temperature overnight. The leaves were removed and decolorized in chloral decolorizing solution for three days, and the decolorizing solution was changed once a day. After decolorization, the spores and mycelium were observed with super field depth stereomicroscope. 100 mL of trypan blue staining solution was formulated with 50 mg of trypan blue, 25 mL of lactic acid, 23 mL of water-soluble phenol (7%), 25 mL of glycerol, and 27 mL of water. The decolorizing solution was 100% chloral solution (Zhang et al. 2018).

### subcellular localization

The cds sequences of WRKY40 and WRKY1 were constructed on pCAMBIA1302 vector, respectively, and the recombinant plasmids transformed EHA105 receptor cell. Summer is the rapid growth period of tobacco, at this time, 5 fully expanded leaves of robust tobacco were selected for Agrobacterium infestation test, the photoperiod was 16h light and 8h dark, incubation for 2 days at 30 degrees. Laser confocal apparatus was used for subcellular localization analysis. DAPI was used as a dye for nuclear localization (Yao et al. 2019).

### Conservative structural domain analysis

The gene sequences of Arabidopsis thaliana were obtained from the TAIR website and the apple genome files were obtained from the GDR website (Daccord et al. 2017). After obtaining the protein sequences of the Arabidopsis genes, the comparison to the apple genome was done to obtain the gene IDs of the apples, and the comparison was done through the Blast plugin of TBtools (Chen et al. 2020). Conserved structural domain analysis was accomplished through the Batch CD-search tool on the NCBI website.

## Author Contributions

Liming Lan and Sanhong Wang designed the research and wrote the manuscript; Liming Lan, Lifang Cao, Lulu Zhang, and Weihong Fu performed experiments for the research and analyzed the data. Shenchun Qu and Sanhong Wang gave the project discussion and instructions. All authors read and approved of its content.

## Acknowledgments

The authors would like to thank Sanhong Wang for helpful discussions on topics related to this work. The authors are also grateful to Lifang Cao for her help with the preparation of figures in this paper.

